# AFM images of open and collapsed states of yeast condensin suggest a scrunching model for DNA loop extrusion

**DOI:** 10.1101/2019.12.13.867358

**Authors:** Je-Kyung Ryu, Allard J. Katan, Eli O. van der Sluis, Thomas Wisse, Ralph de Groot, Christian Haering, Cees Dekker

## Abstract

Structural Maintenance of Chromosome (SMC) protein complexes are the key organizers of the spatiotemporal structure of chromosomes. The condensin SMC complex, which compacts DNA during mitosis, was recently shown to be a molecular motor that extrudes large loops of DNA. The mechanism of this unique motor, which takes large steps along DNA at low ATP consumption, remains elusive however. Here, we use Atomic Force Microscopy (AFM) to visualize the structure of yeast condensin and condensin-DNA complexes. Condensin is found to exhibit mainly open ‘O’ shapes and collapsed ‘B’ shapes, and it cycles dynamically between these two states over time. Condensin binds double-stranded DNA via a HEAT subunit and, surprisingly, also via the hinge domain. On extruded DNA loops, we observe a single condensin complex at the loop stem, where the neck size of the DNA loop correlates with the width of the condensin complex. Our results suggest that condensin extrudes DNA by a fast cyclic switching of its conformation between O and B shapes, consistent with a scrunching model.

## INTRODUCTION

The structural maintenance of chromosomes (SMC) family of protein complexes, such as condensin, cohesin, and the SMC5/6 complex, is pivotal to cellular life (Aragon et al., 2013; Hirano, 2016; Jeppsson et al., 2014; Nasmyth and Haering, 2005; Rowley and Corces, 2018; van Ruiten et al., 2018; Uhlmann, 2016). These SMC protein complexes play vital roles for spatially organizing chromosomes as they are key in sister chromatid cohesion, chromosome condensation and segregation, DNA replication, DNA damage repair, and chromosome resolution (Haering and Gruber, 2016; Nasmyth and Haering, 2005). Condensin plays a crucial role to compact mitotic chromosomes during cell division (Hirano, 2016). Indeed, when condensin is defective, both chromosome formation and the resolution of sister chromatids fail (Houlard et al., 2015; Oliveira et al., 2005). Structurally, SMC protein complexes feature a circularly connected conformation of a kleisin–SMC heterodimer ring (Figure 1A): Two anti-parallelly folded coiled-coil arms, Smc2 and Smc4 (each ~45 nm in length), feature an interconnected hinge domain at one end and heads with an ABC-type nucleotide-binding domain at the other end, where the two head ends are mutually connected by a Brn1 kleisin. Furthermore, two HEAT-repeat subunits, Ycg1 and Ycs4, are bound to the kleisin. A recent crystal structure showed that DNA is anchored by Ycg1–Brn1 through a safety-belt mechanism (Kschonsak et al., 2017).

**Figure 1.**
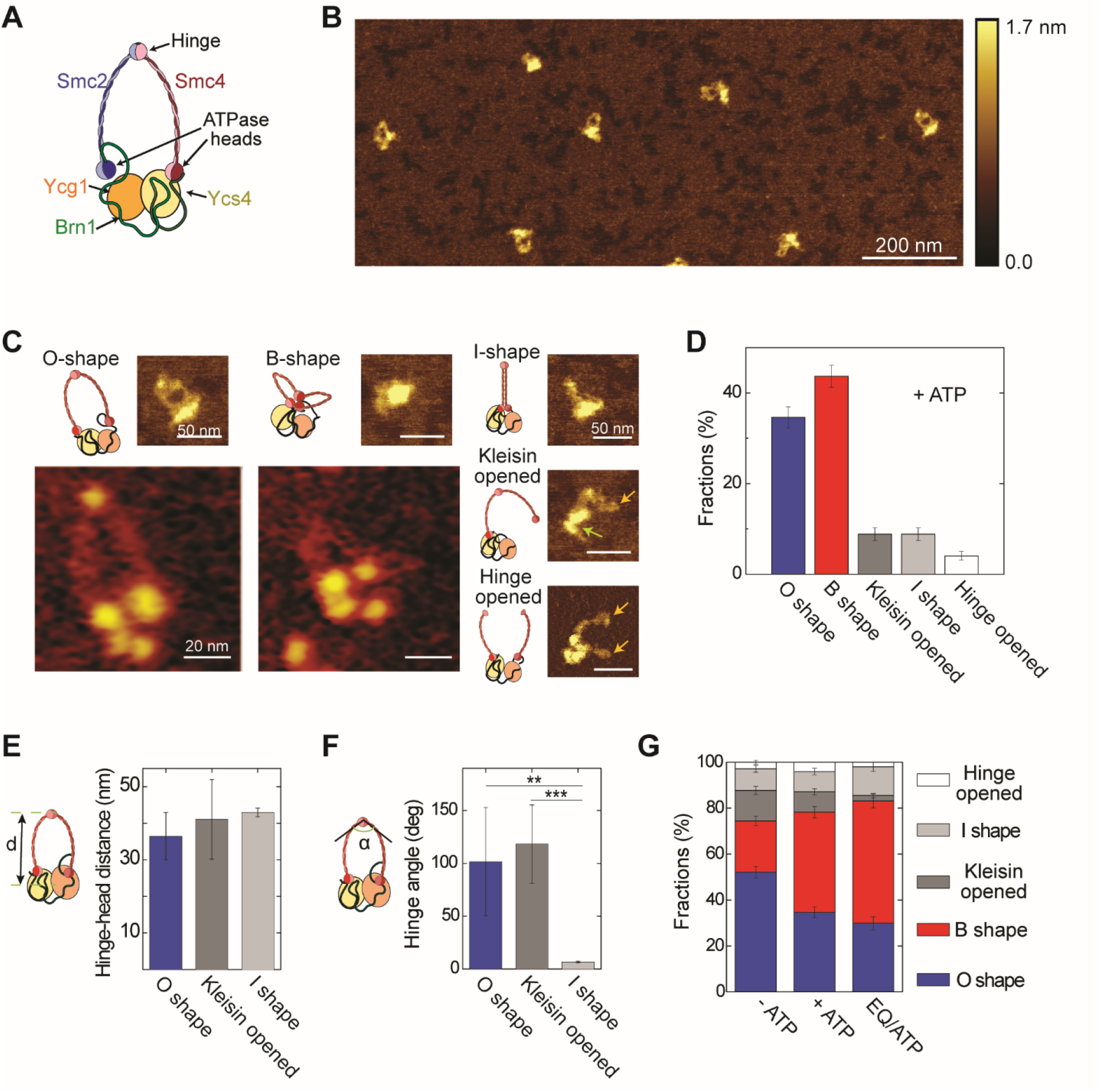
The condensin holocomplex exhibits different shapes. (A) Cartoon of the yeast condensin holocomplex. (B) Typical dry AFM image of condensin. (C) Images of various conformations of condensin holocomplexes (*N* = 384). Small panels are dry-AFM images and large panels are liquid-AFM images of the O- and B-shape. In the AFM image of kleisin-opened state, the green arrow indicates the non-SMC domains while the orange arrow points to an smc end. In the AFM image of the hinge-opened state, orange arrows indicate the dissociated hinge domains. (D) Relative occurrence of different states in the presence of ATP. (E) Distance between hinge and the globular domain for O shape, kleisin-opened shape, and I-shape condensins. (F) Hinge angle measured at the hinge domain. * *P*<0.05, ** *P*<0.01, and *** *P*<0.001, assessed using the paired *t* test. (G) Relative occurrence of various conformations (*N*=384 without ATP, *N*=255 with ATP, and *N*=419 for the EQ mutant with ATP). Error bars indicate SD unless specified otherwise.

Hi-C studies have suggested that mitotic chromosomes consist of DNA loops extruded by SMC protein complexes (Dekker and Mirny, 2016; Fudenberg et al., 2016; Naumova et al., 2013; Rao et al., 2017; Sanborn et al., 2015) In addition, polymer simulations indicated that loop extrusion can result in the efficient disentanglement and compaction of chromatin fibers (Alipour and Marko, 2012; Goloborodko et al., 2016). The formation of these loops, which can be very long (up to Mbp), requires a processive motor that extrudes DNA. Recent single-molecule fluorescence studies indeed demonstrated a motor function (Terakawa et al., 2017) and provided direct evidence for DNA loop extrusion by the yeast condensin complex (Ganji et al., 2018). Condensin constitutes a unique new type of DNA-translocase motor that reels in DNA at high speed (up to 1.5 kbp/s at low forces) while consuming low ATP, i.e. an ATPase rate of ~2 ATP/s, implying that the motor takes very large (~50 nm) steps per ATP molecule that is hydrolyzed (Ganji et al., 2018). While this large step size suggests that the motor action involves sizeable conformational changes at the level of the size of the full SMC complex, it is currently unclear by what molecular mechanism condensin is able to process DNA so efficiently. Various models have been suggested (Hassler et al., 2018). For example, in a tethered inchworm model (Nichols and Corces, 2018), condensin moves along DNA by a sequential binding of the heads to DNA, much like the motion of kinesin along microtubules. In a DNA-pumping model (Diebold-Durand et al., 2017; Marko et al., 2019), a zippering of two SMC coiled-coil arms pushes entrapped DNA from the hinge domain to the head domains, while an ATP-induced dimerization of the head domains changes the condensin conformation from an I-shape to O-shape to target new binding of DNA for subsequent cycles. In a scrunching model (Terakawa et al., 2017), an extension and retraction of the flexible SMC arms move DNA connected to the hinge domain to the globular subunits near the heads, and back in a cyclic fashion. Although the open question on the motor mechanism is at the heart of resolving the functioning of SMC proteins complexes, and hence is crucial for understanding chromosomal structure, experimental evidence for any of these models has thus far been very limited.

Studying the conformational changes of condensin and its connection to the mechanochemical cycle that couples ATP hydrolysis to structural changes of the protein complex is thus critical to the understanding of the mechanism underlying DNA loop extrusion by condensin. Many studies have been undertaken to resolve the structure of a variety of SMC proteins, with contradictory results so far, however (Eeftens and Dekker, 2017). Negative-staining EM structures and crosslinking experiments suggested a rigid extended structure (I-shape) of prokaryotic SMCs, while ATP-dependence studies indicated a transition from I- to O-shape upon ATP hydrolysis (Diebold-Durand et al., 2017; Kamada et al., 2017; Soh et al., 2015). A recent EM and cross-linking study on MukBEF and yeast cohesin suggested transitions between an I shape and a folded state (Bürmann et al., 2019), similar to an early AFM study on *S. pombe* condensin (Yoshimura et al., 2002). High-speed (HS) atomic force microscopy (AFM) images of yeast condensin SMC dimers (without the kleisin and HEAT-repeat co-factors) in liquid showed instead very flexible SMC arms and dynamical conformational changes between O-, V-, P-, and B-shapes (Eeftens et al., 2016). However, thus far, no HS liquid AFM studies on the full condensin holocomplex have been reported. High-resolution imaging of the yeast condensin holocomplex with all its subunits is highly desirable in order to understand its conformations at the stem of the DNA loop, where it reels in DNA.

Here, we resolve the conformational states and shape transitions of yeast condensin and its binding configuration to DNA, with the aim to understand the motor dynamics that drive DNA loop extrusion. We use both dry AFM and liquid HS AFM for visualization. AFM imaging comes with significant advantages: it features a high resolution (xy resolution ~ 1 nm, z resolution ~ 0.1 nm), it provides high contrast, and it is label free and cross-link free, making it very suitable to study complex DNA-protein interactions. Dry AFM imaging, where the sample is dried after incubating the molecules on a surface, enables high-throughput imaging by capturing many molecules in various conformations, including intermediates of loop extrusion. HS AFM in liquid even enables imaging of real-time structural changes of condensin within a physiological buffer in the presence of ATP, with a temporal resolution up to 20 frames per second, although it features a lower throughput due to small imaging area (~100×100 nm^2^) that is needed to realize a high frame rate (Katan and Dekker, 2011; Kodera et al., 2010; Uchihashi et al., 2011).

A major finding from our data is that the condensin holocomplex predominantly exhibits either an open O-shape or a collapsed B-shape, and that it cycles back and forth between these states. In addition, we find that the SMC arms are highly flexible and condensin binds to DNA through both the hinge and HEAT-repeat subunits. This is suggestive of a scrunching model for this DNA-loop-extruding motor, which involves a cycle where DNA is periodically transferred between two different DNA-binding sites, near the hinge and near the globular HEAT/head subunits.

## RESULTS

### Condensin exhibits an open O-shape or collapsed B-shape

Figure 1B shows an example of an AFM image of individual *Saccharomyces cerevisiae* condensin holocomplexes, i.e., the full complex with all its 5 subunits. These purified proteins showed DNA-stimulated ATPase activity and DNA loop extrusion (Figure S1), similar to earlier reports (Ganji et al., 2018). The AFM images showed a homogeneous volume distribution of the condensins (Figures 1B and S2A), with an average complex volume of 2,405 ± 430 nm^3^ (*N* = 384). Similar to previous AFM studies that resolved the structure and various conformational states of the arms of the ‘SMC dimer only’ Smc2–Smc4 complexes, we were able to resolve the SMC arms in most of our condensin images, as well as distinguish various conformational states of the condensin holocomplexes (Figures 1C-G).

The main configurations that condensin was found to adopt were O-shapes and B-shapes (Figures 1C and D). In the O-shape, a ring-like configuration was clearly identified, with two flexible arms, a large globular domain at one side, and a locally increased height at the hinge at the other side. The globular domain is presumably composed of the two SMC head domains, two HEAT-repeat subunits (Ycg1 and Ycs4), and the Brn1 kleisin subunit. In dry AFM, these could not be resolved. In contrast, using liquid-phase AFM, we resolved multiple individual subunits (Figure 1C): Smc2–Smc4 head domains and two HEAT-repeat subunits. The arms were seen to adopt many different configurations, demonstrating a high flexibility. The O-shapes exhibited on average a distance of 37 ± 7 nm (mean ± s.d.) between the center of mass of the globular and hinge domains (Figure 1E), and featured a 102 ± 51 degrees opening angle at the hinge (Figure 1F). The second prominent state of condensin that we observed was a collapsed B-shape. This state appears to correspond to a ‘butterfly shape’ formed when the hinge domain engages the joined SMC head domains, similar to observations in SMC-dimer-only samples (Eeftens et al., 2016). To check whether surface charge affects the conformation of the proteins, we also imaged complexes on polylysine-treated mica, which is a positively charged surface, instead of on mica, which is slightly negatively charged (as in the data in Figure 1). This yielded the same result (Figure S3), viz., a dominance of O shapes and B shapes, indicating that these conformations are not induced by electrostatic surface-protein interactions.

Together, the O shapes and B shapes accounted for a high fraction (~80 %) of all data. Minority states that were observed were I-shapes (9 %), kleisin-opened shapes (9 %), and hinge-opened states (4 %). In the I-shapes, the two SMC arms are co-aligned, and the distance between hinge and non-SMC subunits was slightly larger than the distance of O-shapes (Figure 1E). Finally, we observed condensins with a disconnection between one of the SMC heads and the non-SMC subunits (indicating a ‘kleisin-opened shape’; Figure 1C), and shapes where the two SMC arms were disengaged at the hinge (‘hinge-opened shapes’). These, relatively rare, states where the complex is partially opened may possibly relate to a topological loading of condensin onto DNA, for which the ring has to open.

Different ATP states yielded a different distribution of O- and B-shapes. We performed AFM experiments under several conditions: with ATP (*N* = 255), without ATP (*N* = 384), and using an EQ mutant of condensin that binds ATP but is unable to hydrolyze it (*N* = 419). In the presence of ATP (i.e., the data discussed above), there were slightly more B-shapes than O-shapes. In contrast, significantly more open O-states were observed without ATP, while instead more collapsed B-shapes were observed for the EQ mutant with ATP (Figure 1G). The latter suggests that the collapsed B state may be promoted by ATP binding to the head domains, whereas the observation that O states were more abundant without ATP may indicate that hydrolysis and release of ATP are associated with a release of the hinge domain from the head domains.

### Condensin cycles back and forth between an open O-shape and collapsed B-shape

We applied HS AFM imaging to visualize the dynamics of the conformational changes of condensin holocomplexes in a buffer with 50 mM NaCl and ATP – equivalent to the buffer where condensin exhibits loop extrusion (Figure S1C; Ganji et al., 2018). Movies were recorded at 5 frames per second (Figure 2; Movies S1–S4). We could resolve 3 or 4 substructures within the globular domain, see e.g., Figure 2A and S4, where one can distinguish, next to the SMC arms and hinge domain, the dimerized head domains and two HEAT-repeat subunits. The SMC arms in the condensin holocomplex were found to be very dynamic and flexible, changing conformations very fast (i.e., at a <0.2 s time scale in this example of a 5-Hz movie, Figure 2A), very similar to previous studies on SMC-dimer-only samples (Eeftens et al., 2016). For the O-shapes, we also observed that the hinge angle between the two SMC arms changed dynamically (Figure S6), consistent with large conformational changes in the hinge domain.

**Figure 2.**
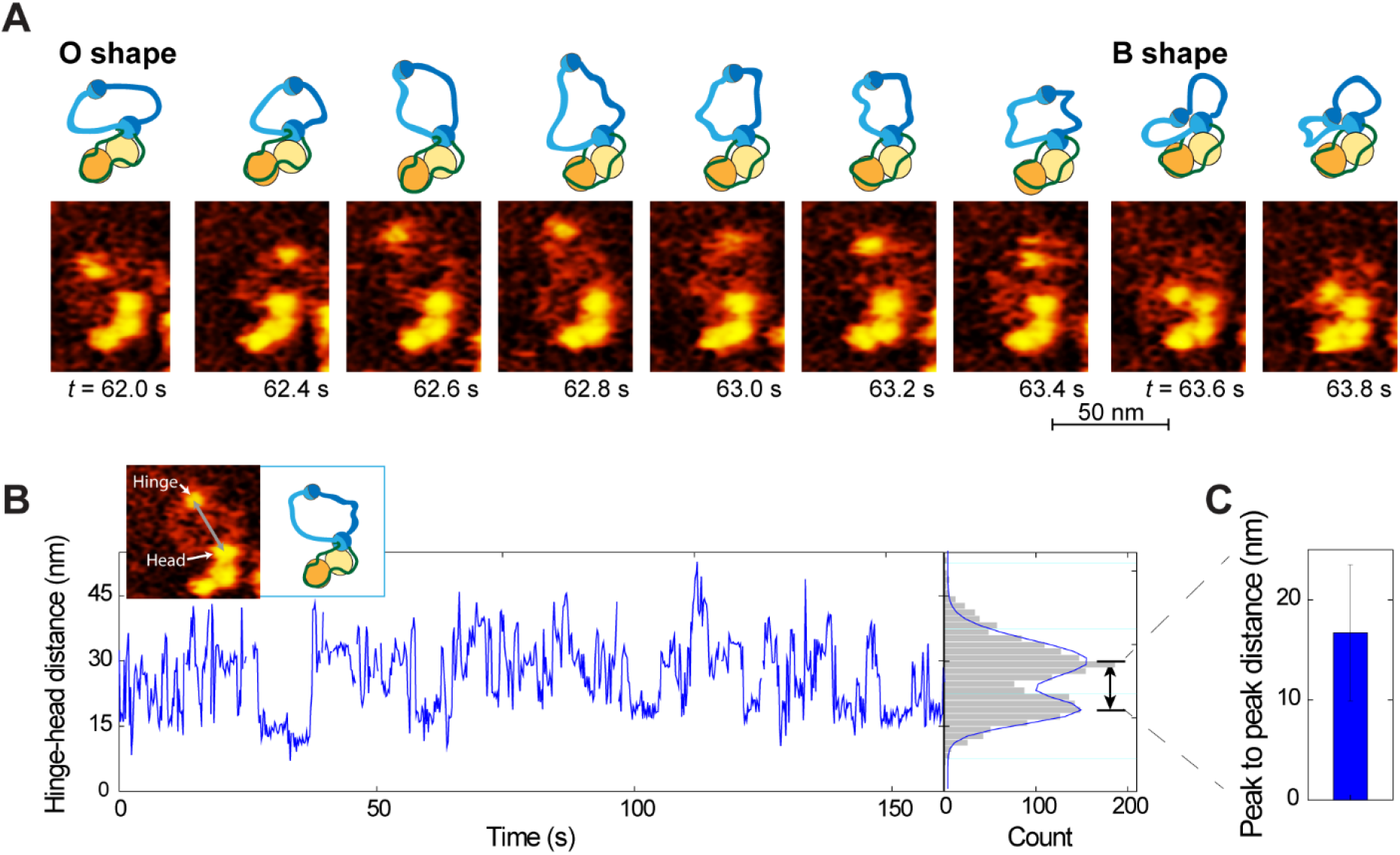
Condensin dynamically toggles between an O-shape and B-shape over time. (A) Representative HS liquid AFM images of the condensin holocomplex followed over time at a 5 frames/s rate (Movie S1). The complex is seen to change its conformation over time, from an O shape (62-63.5 s) to a collapsed B-shape (after 63.5-64 s). (B) Real-time trace of the hinge-head distance. The distance is seen to toggle between a large and small value, corresponding to the O-and B-shapes respectively. The distribution of the hinge-head distance, depicted on the right, clearly shows two peaks. Solid line is a fit of two Gaussians. (C) Average peak-to-peak distance of 17 ± 7 nm from many distributions such as in panel B (*N*=12 individual traces taken on different SMC molecules).

Interestingly, we observed clear dynamic conformational changes where condensin would toggle between an O-shape and a collapsed B-shape, and vice versa (Figures 2A-2B, and S4; Movie S1-4). In these movies, the hinge region transitioned between a far-away position and a location close to the globular domain, consistent with an overall shape change between the O-and B-states, respectively (Figures 2A-2B). Quantification of this dynamics is provided in Figure 2C, which plots the hinge-head distance as measured between their centers of masses (cf. inset to Figure 2B). The data clearly exhibit a two-level-fluctuator behavior, displaying two clear peaks in the histogram (right side of Figure 2B). The distance between these peaks, averaged over multiple such traces (*N* = 12), was 14 ± 4 nm (mean ± s.d; Figure 2C). The transition between the O-shape and the B-shape state can be completed very fast, faster than our temporal resolution (0.2 seconds in this movie). Note that the hinge movement was not strongly affected by tip-sample interaction, because, as shown in Figure S5, we find that the hinge movement is isotropic and not biased by the AFM scanning direction, indicating that the hinge displacement rather reflects the intrinsic behavior of this domain.

### Condensin binds double-stranded DNA via a HEAT subunit and via the hinge domain

In order to reel in DNA to extrude a DNA loop, the condensin motor complex should exhibit multiple binding sites to DNA. Identification of the DNA binding sites is therefore important. We used dry AFM to examine the interaction between condensin and DNA. We imaged large scan areas (>10 ×10 μm^2^) in order to gather good statistics in counting bound condensin complexes. The AFM images showed different binding modes (Figure 3). First, as expected, we observed that condensin binds to DNA at the globular domain (Figure 3A), presumably at the Ycg1–Brn1 interface (Kschonsak et al., 2017). Second, we observed binding of condensin to DNA via the hinge domain (Figure 3B). Unexpectedly, hinge binding was observed to occur very frequently, almost as often as HEAT binding, viz., for 35 versus 33 % of the complexes, respectively (Figure 3E). Third, we observed that condensin can bind DNA through both the hinge and globular domains (Figure 3C). Condensin was also observed to bind to the DNA in the collapsed B-state (26%; Figure 3D), but here it was difficult to discriminate the exact binding location within the complex.

**Figure 3.**
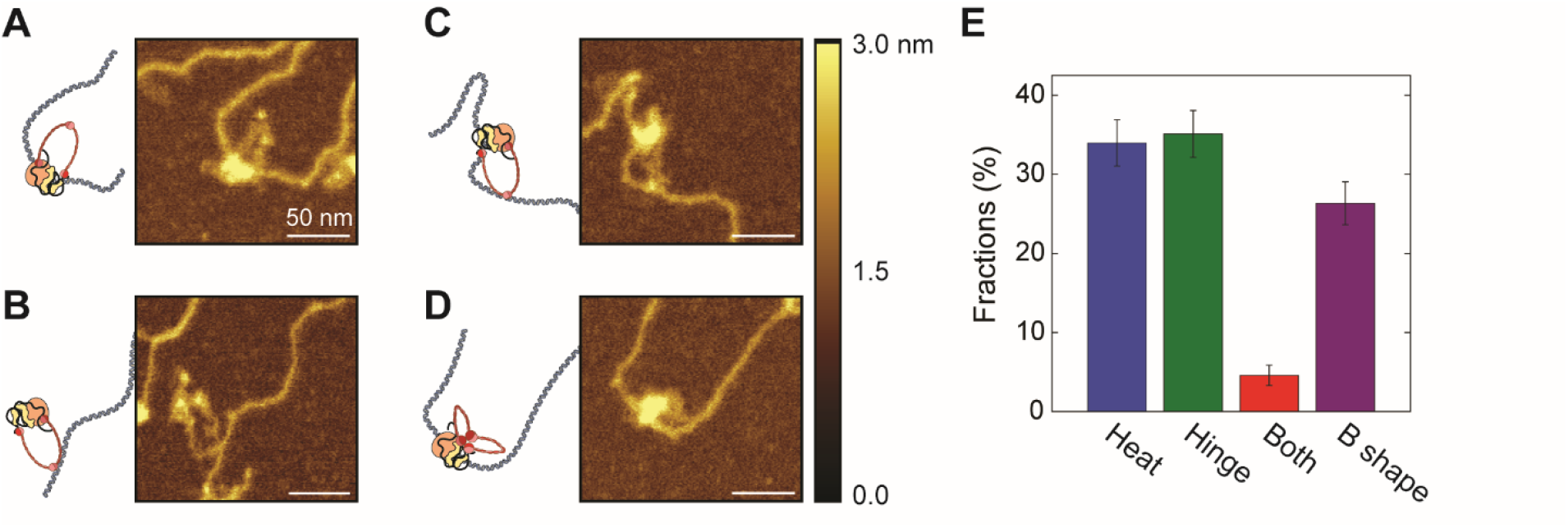
Binding of condensin to DNA. (A) Binding of condensin to DNA through its globular domain, presumably through the Heat domains. (B) Binding of condensin to DNA through its hinge. (C) Binding of condensin to DNA at two sites. (D) Collapsed B-shape condensin on DNA. (E) Relative occurrence of each of the different binding modes (*N*=262).

A number of controls supported these results. Firstly, we established that an accidental colocalization, i.e. a possible fortuitous overlap of DNA and unbound condensin, contributes only minor fraction in the presence of ATP (Figure S7), indicating that condensin binding to DNA is driven by biochemical interactions. Secondly, the DNA-binding probability of wild-type condensin was much higher than that for a complex that lacks the Ycg1 subunit (Figure S7B), consistent with a previous study that Ycg1 is important for DNA binding (Piazza et al., 2014). Finally, we observed a decrease in the number of condensins binding to DNA in experiments where we omitted ATP or replaced it with ATPγS (Figure S7B), consistent with previous observations that ATP hydrolysis is needed to topologically load condensin onto DNA (Eeftens et al., 2017) and increase condensin’s residence time on DNA (Ganji et al., 2018).

### DNA loops feature a single condensin complex at their stem

Next, we used AFM to image condensin holocomplexes at the stem of loops of DNA that were formed by condensin. To observe such extruded loops in dry AFM images, it is important to avoid stretching forces on the DNA during the drying process, since forces as low as ~10 pN are enough to disrupt the loops formed by condensin (Eeftens et al., 2016). Using a careful drying procedure (see Materials and methods), we could image condensins on extruded loops of DNA with dry AFM. To prepare samples, we mixed condensin with DNA and ATP in a test tube and incubated the mixture for 3 min, an incubation time similar to the lag time between condensin arrival and the initiation of DNA compaction observed in a magnetic tweezers experiment (Eeftens et al., 2017). For dry AFM imaging, we subsequently deposited the samples under a very slight stretching force onto a mica surface that was pre-functionalized with a low concentration of polylysine, while for liquid-phase AFM, we immobilized the sample onto mica with the addition of 0.3 mM spermidine.

Both in liquid and dry AFM, we observed a clear co-localization of condensin and the stem of DNA loops (Figures 4A-B; Movie S5). We conclude that these loops were predominantly formed by condensin in an ATP-hydrolysis-dependent manner, based on the following observations. First, the number of co-localizations was much higher than could be expected from random colocalization (Materials and Methods; Figures S8). Second, the loop size of DNA was found to be significantly larger for the case of incubation with ATP than for any negative controls (Figures 4E and S9). Moreover, the number of condensin-mediated loops depended strongly on the availability of hydrolysable ATP (Figure 4C). Negative controls with ATPγS or without nucleotide showed a significantly lower number of loops per unit DNA length, 0.029 loops/μm and 0.019 loops/μm respectively, compared to 0.15 loops/μm for the case with ATP (counting, in this case, only loops that have a condensin at their stem; see also Figure S8D). And finally, we rarely observed loops when using a tetrameric a *S. cerevisiae* condensin complex that lacks Ycg1 (Figure 4C). Taken together, these experiments show that condensin localizes to the loop stem when extruding DNA loops.

**Figure 4.**
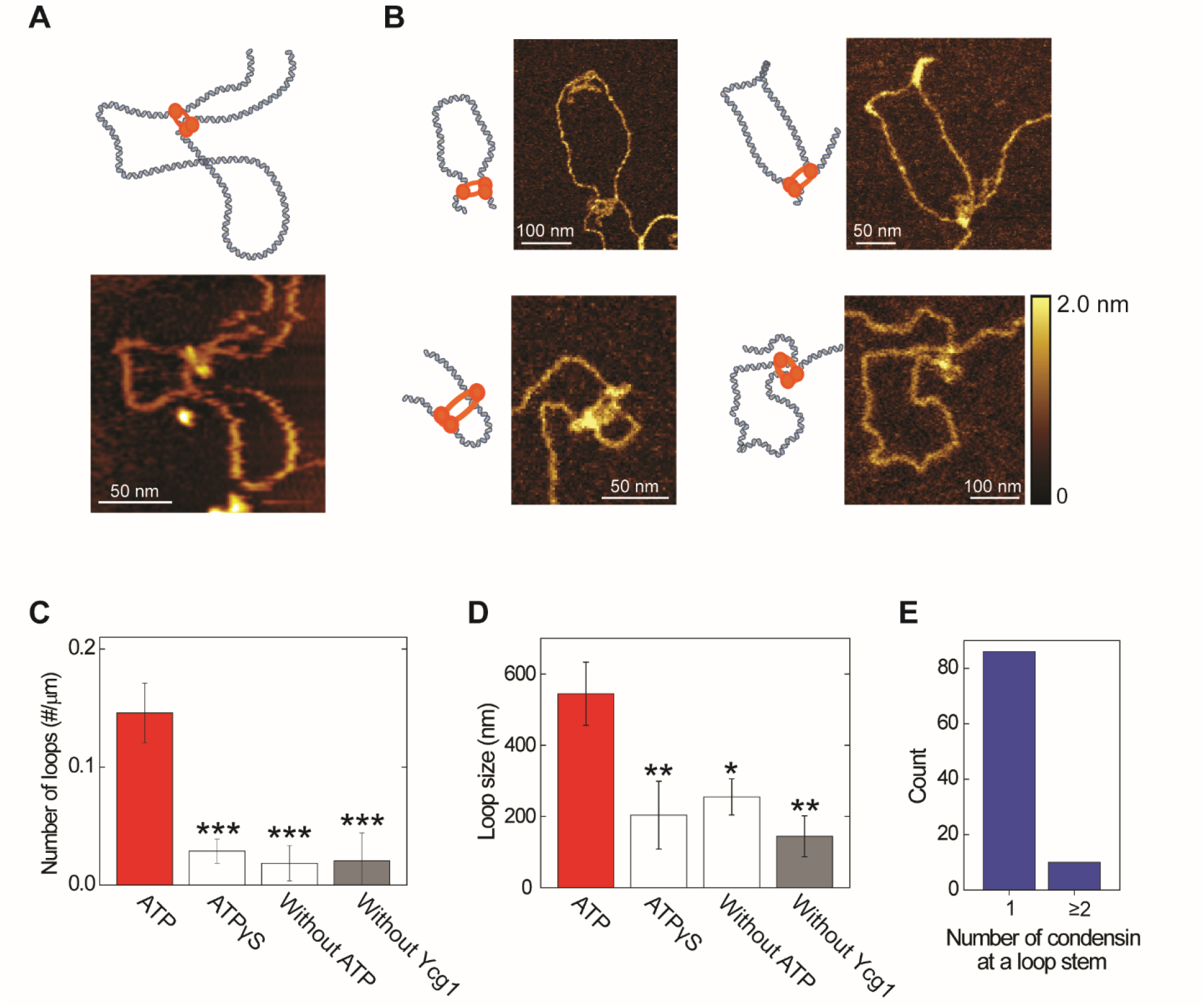
Condensin-mediated DNA loops. (A-B) Representative condensin-mediated DNA loops imaged using liquid AFM (A) and dry AFM (B). Cartoons are a guide to the eye. (C) Number of DNA loops per μm of DNA under various conditions: with ATP (150 loops), with ATPγS (21 loops), without ATP (24 loops), and without Ycg1 (15 loops), as measured on *N* ≥5 independent experiments. Loop formation is strongly dependent on ATP hydrolysis and on the presence of the Ycg1 domain. (D) Loop size comparison in various conditions (median ± SEM). (E) Distribution of the number of condensins found at the stem of a DNA loop (*N=*96).

We quantified the number of condensin molecules at the stem of the DNA loops by estimating their volume. We found that 90 % of the DNA loops showed a single condensin at the loop stem (Figure 4E), consistent with an earlier estimate from optical measurements (Ganji et al., 2018) and confirming that a single condensin holocomplex is responsible for loop extrusion of the DNA. We rarely (10%) observed multimeric forms of condensin at the stem of the DNA loops in our experimental conditions. Our observation of single condensins at the DNA loop stem contrasts suggestions of the handcuff model where loop extrusion by cohesin or MukBEF was suggested to be driven by dimeric SMC complexes (Woo et al., 2009; Zhang et al., 2008).

### The neck width of the DNA loop stem correlates to the size of the condensin complex

Next, we examined the conformation of protein and DNA at the loop stem in more detail. Specifically, we examined the correlation between the neck width at the stem of the DNA loop (Figure 5) and the width of the condensin molecule located there. The neck width of the DNA loop was determined from the distance between the two pieces of DNA right next to the condensin at the loop stem, as measured from the height profile of the cross section across the two pieces (Figure 5A, bottom). The condensin width was similarly obtained from the height profile of the cross section. For good comparison, both the DNA neck width and protein width were defined as the distances between two outer points of the full width half maximums of the Gaussian fits, since separated peaks could not always be resolved when the two DNA molecules were too close.

**Figure 5.**
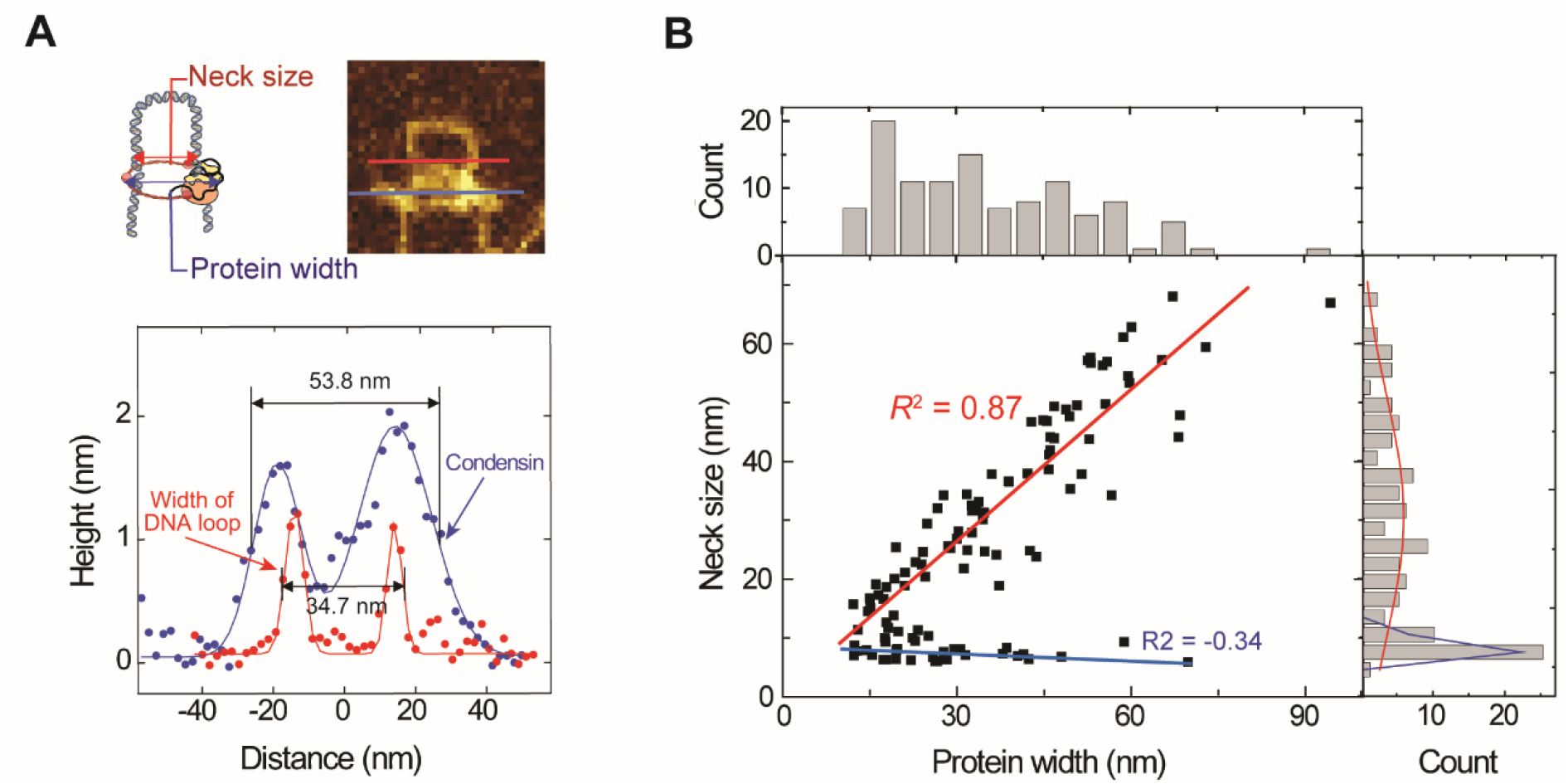
The neck size of the DNA loop correlates with the size of the condensin complex. (A) Top left schematic indicates the protein width and neck size of the DNA loop. Lines in the image on the top right indicate where crosssectional height profiles are acquired, with the result denoted at the bottom panel. (B) Neck size of DNA loops versus width of the condensin at the loop stem. Every point comes from a measurement of a single loop (*N*=115). The distribution of neck sizes (shown on the right) is fitted to two Gaussian, yielding average peak values of 9 ± 1 and 31 ± 20 nm.

Interestingly, we found a clear linear relationship between the width of the DNA loop at the stem and the characteristic size of the protein, as measured at the neck (correlation coefficient *R*^2^ = 0.87; Figures 5B and S10C). This strong tendency to follow the protein conformation suggests that DNA is bound to a condensin complex that varies in size. Next to the population that showed this linear correlation, a second smaller population (blue data in Figure 5B; and S10D) showed no dependence of the DNA neck width on condensin size. From studying the individual images (Figure S10), we conclude that the small DNA-neck width of these molecules is caused by interwound structures within the DNA, that likely are induced upon depositing the DNA onto the polylysine-treated mica surface (Japaridze et al., 2017a). Overall, the data suggest that conformational changes of the condensin complex appear to be associated with changes in the width of the stem of the DNA loop that is extruded by condensin.

## DISCUSSION

Summing up, we visualized yeast condensin holocomplexes on their own as well as bound to DNA. We found that condensin predominantly exhibits an O-shape or collapsed B-shape, and that it dynamically switches between these conformations. When condensin was added to DNA, it bound at the hinge or at the globular domain that is formed by the SMC heads and HEAT-repeat subunits. Condensin formed DNA loops, and we observed that a single condensin was located at the loop stem, where the loop-stem width correlated with the condensin size. Below we discuss the relevance and importance of these findings.

### Flexible SMC arms

Our liquid HS-AFM imaging showed very flexible SMC arms with highly dynamic conformational changes. This is very similar to previous observations for coiled coil arms of the SMC dimers only, which exhibited a persistence length of only 4 nm (Eeftens et al., 2016), indicating that the flexibility of SMC arms is a general feature of the condensin holocomplex and not influenced by the interaction between SMC arms and non-SMC subunits. While the HS-AFM images unambiguously show flexible SMC arms, more rigid SMC arms were implied in reports using other techniques, such as X-ray crystallography, cross-linking experiments, or EM (Diebold-Durand et al., 2017; Soh et al., 2015). For Rad50, a related SMC-like protein, liquid phase imaging similarly showed flexible and dynamical SMC arms (Anderson et al., 2001; Moreno-Herrero et al., 2005), while crystal structures showed rigid rods (Park et al., 2017). HS-AFM thus appears to be a liquid-phase imaging technique that can visualize dynamical changes of the SMC complexes with high spatial and temporal resolution, which may be missed in X-ray crystallography, cross-linking experiments, and EM imaging of fixed samples.

### Conformation of the condensin complex

Our data show that condensin predominantly exhibits an extended O-shape or collapsed B-shape. This fits observations in an early AFM study and a recent EM paper that showed both extended and compacted configurations of SMC proteins (Yoshimura et al., 2002; Bürmann et al., 2019). However, our data did not show a juxtaposed I-shape that was reported in these and other studies (Soh et al., 2015; Diebold-Durand et al., 2017). There are several possible explanations for the discrepancy of O- versus I-shape observations among various studies. First, SMC complexes from different organisms may exhibit different shapes. For example, *S. cerevisiae* cohesin and non-crosslinked MukBEF were found to adopt a folded conformation (Bürmann et al., 2019), while *B. subtilis* SMC proteins showed I-shapes and O-shapes (Diebold-Durand et al., 2017). Second, the condensin complex is a large protein complex and purifying intact functional holocomplexes is challenging. We verified that our proteins were functional and capable of loop extrusion (Figure 4 and S1). Third, different sample-preparation methods can induce or bias the observation of certain configurations. The high signal-to-noise ratio in AFM imaging enables to analyze every single molecule in the field of view, which prevents the problem of selection bias that is an ever-luring danger in single-particle electron microscopy (Lyumkis, 2019). Furthermore, the air-water interface in cryo-EM samples can cause a preferred orientation of proteins (Noble et al., 2018) or denature proteins (D’Imprima et al., 2019). The sample preparation for AFM is relatively mild and no staining or labeling is required. From the AFM imaging in solution, it is clear that the coiled-coil arms and hinge domain are parts of the complex that interact only weakly with the surface, which facilitates the condensin to adopt an equilibrated shape. However, the presence of a surface is a requirement in AFM (as also is the case in most EM techniques), which potentially may affect the condensin structure. We measured both on slightly negatively charged mica surfaces and on positively charged polylysine-treated surfaces (Figure S3), and O-shapes and B-shapes were the most abundant in both cases, suggesting that these shapes are intrinsic to condensin.

### DNA binding sites of the condensin complex

Our AFM images of DNA-bound condensin show that condensin can bind DNA at (at least) two binding sites, one at the hinge and one at the globular domain with the SMC heads and HEAT-repeat subunits. Previous work clearly showed that Ycg1-Brn1 provides a strong DNA binding site (Kschonsak et al., 2017), which is consistent with the binding that we observed near the globular domain that includes the HEAT-repeat subunits (Figure 3A). The frequent occurrence of hinge binding that we observed is more surprising. Since the inner region of the hinge domain has a positively charged surface, it is actually plausible that this may bind DNA (Griese et al., 2010). Indeed, gel-shift assays and fluorescence anisotropy experiments showed DNA binding affinity to the hinge domain of mouse condensin (Griese et al., 2010), yeast condensin (Piazza et al., 2014), bovine cohesin (Chiu et al., 2004) and prokariotic condensin (Hirano, 2016). Previous AFM studies also showed DNA binding by the hinge domain for MukB (Kumar et al., 2017) and for *S. pombe* condensin (Yoshimura et al., 2002), suggesting a conserved hinge-binding mechanism. Our observation that the hinge angle of the O-shaped condensin changes dynamically (Figure S6) indicates that the hinge domain is very flexible. This is consistent with previous crystal structures of hinge domains that showed two different states of the fully associated and half-dissociated conformations (Griese et al., 2010; Haering et al., 2002; Ku et al., 2010; Li et al., 2010; Niki, 2017).

### Implication for models of DNA loop extrusion by SMC complexes

Given that it thus far has remained puzzling how the condensin complex is able to extrude a DNA loop, it is of interest to discuss what our findings imply for a mechanistic model for SMC-induced DNA loop extrusion and to see how it relates to various models that have been proposed. A working model should account for a number of specific experimental observations: it should (i) involve cyclic conformational transitions between an O-shape and B-shape of the SMC complex, (ii) be compatible with very flexible SMC arms, (iii) accommodate a very large step size (Ganji et al., 2018), (iv) show a high correlation between the condensin size and the neck width at the stem of DNA loops (Figure 5), (v) use Ycg1-Brn1 as an DNA anchoring site (Ganji et al., 2018; Kschonsak et al., 2017), and (vi) use a DNA-binding site at the hinge domain.

Our experimental results appear to rule out the ‘tethered inchworm model’ (Nichols and Corces, 2018), which predicts that condensin moves along DNA by a sequential binding of the heads, similar to the motion of kinesin along microtubules. Instead of the prominent head-head motions involved in this model, we observed substantial hinge-head motions. Furthermore, this model assumes that the hinge domain anchors the DNA, which is contradictory to the finding that Ycg1-Brn1 is a DNA anchoring site. Furthermore, the ‘DNA pumping model’(Diebold-Durand et al., 2017) is also inconsistent with our observations. In the DNA pumping model, a zipping of the SMC arms from an O- to I-shape pushes DNA from the hinge to the head domains, whereupon a subsequent conformational change from I- to O-shape targets new DNA. The I-shapes thus play a prominent role in this model, whereas a B-shape does not occur in this cycle – both in stark contrast to our observations.

Our data instead indicate that some form of a ‘scrunching model’ underlies the motor action of SMCs (Figure 6), where condensin anchors to DNA and moves another piece of the DNA relative to this point to extrude a loop, by cyclically binding DNA to the hinge and reeling it to the globular heads/non-SMC domain. In such a model, the hinge binds DNA, transfers it to the head region, binds it there, upon which the hinge releases to subsequently grab a new piece of DNA to reel it in (Terakawa et al., 2017; Hassler et al., 2018). Our observation that condensin dynamically switches between an O-shape or collapsed B-shape clearly hints at such a type of mechanism. Our finding that the hinge binds DNA further supports this model. The flexible SMC arms provide a hint on the mechanism of DNA transfer from the hinge to the globular domain: While such flexible arms prevent a rigid-body power-stroke motion to directly transduce mechanical motion from the ATPase head domain to the distant (37 nm) hinge domain, the arms instead may use thermal fluctuations to facilitate the motion of the DNA-bound hinge region to the heads/non-SMC subunits as a biased-ratchet motor (Hwang and Karplus, 2019). ATP binding may induce the interaction between the hinge and the head domains to form a B-shape, whereas subsequent ATP hydrolysis may release the hinge (but not the DNA) from the globular domain to restart the cycle. During this cycle, the Ycg1–Brn1 anchor would ensure that condensin remains locked to DNA, such that the newly reeled-in DNA indeed translocates with respect to the anchor position, thus forming a DNA loop.

**Figure 6.**
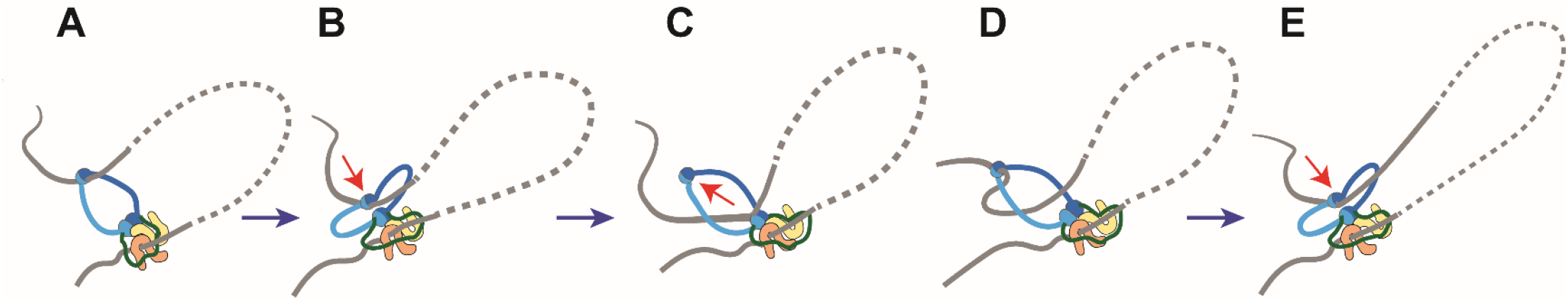
Working model for condensin-mediated DNA loop extrusion. Condensin binds DNA tightly to the Ycg1 anchor, as well as, transiently, to two other binding sites. From an open O-shape (A) it collapses into a B-shape (B) where the neck of the DNA loop concurrently changes from wide to narrow. Subsequently, the hinge is released (C), and the hinge reaches out to bind new DNA (D). (E) Upon repeating this cycle, DNA is extruded into a loop.

Such a scenario does imply the presence of another DNA binding site (next to the hinge and the Ycg1–Brn1 anchoring site) in the globular heads/non-SMC subunits that prevents slipping of the extruded DNA loop while the hinge is released in search of another piece of DNA. Based on previous suggestions for Rad50 and prokaryotic SMC complexes (Erickson, 2009; Liu et al., 2016; Seifert et al., 2016; Woo et al., 2009), such a third DNA-binding site might be located in a positively charged cavity of the ATP-dimerized head domains or in the head-proximal coiled coils (Rojowska et al., 2014). Furthermore, it is still an open question how the unidirectionality of the loop extrusion is established in the scrunching model. This may relate to the angle with which condensin binds DNA at its (transient) binding site, but resolving this will have to wait for further structural studies.

In conclusion, our experimental observations support a scrunching model in which a DNA-bound condensin extrudes a loop of DNA by cycling its conformation between O- and B-shapes. It will be of interest to explore whether such a motor mechanism also applies to other SMC complexes such as cohesin (Davidson et al., 2019; Kim et al., 2019).

## ACKNOWLEDGMENTS

We thank S. Bisht, J. Eeftens, M. Ganji, E. Kim, B. Pradhan, I. Shaltiel, J. van der Torre, and W.W.W. Yang for discussions. This work was supported by ERC grants SynDiv 669598 (to C.D.), Marie Sklodowska-Curie grant agreement No 753002. (to J.-K.R.), CondStruct 681365 (to C.H.H.), the Netherlands Organization for Scientific Research (NWO/OCW) as part of the Frontiers of Nanoscience and Basyc programs, and the European Molecular Biology Laboratory (EMBL).

## AUTHOR CONTRIBUTIONS

J.-K.R., C.D., A.J.K., and C.H.H. designed the experiments, J.-K.R., A.K., and T.W., performed the AFM experiments, E.V.D.S. and J.-K.R. purified condensin complex, J.-K.R., A.J.K., and T.W. contributed image analysis, J.-K.R and R.D.G. performed ATPase assay and single-molecule fluorescence assay, C.D. and C.H.H supervised the work, J.-K.R., C.D., and A.J.K. wrote the manuscript.

## COMPETING INTERESTS

The authors declare no competing interests

## MATERIALS and METHODS

### Condensin holocomplex purification

We used the same protocol as previously reported (Ganji et al., 2018) for the purification of *S. cerevisiae* condensin holocomplexes with all the subunits: Wild-type (Smc2-Smc4-Brn1-Ycs4-Ycg1), Ycg1-missing mutant (Smc2-Smc4-Brn1-Ycs4), and EQ mutant (Smc2_E1113Q_-Smc4_E1352Q_-Brn1-Ycs4). The transformation of these complexes was done using using 2μ-based high copy plasmids containing *pGAL10-YCS4 pGAL1-YCG1 TRP1* (Ycg1-missing mutant) and *pGAL7-SMC4-StrepII3 pGAL10-SMC2 pGAL1-BRN1-His12-HA3 URA3* (wild-type, strain C4491). The overexpressed cells were lysed in buffer A (50 mM TRIS-HCl pH 7.5, 200 mM NaCl, 5% (v/v) glycerol, 5 mM β-mercaptoethanol, 20 mM imidazole) supplemented with 1× complete EDTA-free protease inhibitor mix (11873580001, Roche) in a FreezerMill (Spex), cleared by centrifugation, loaded onto a 5-mL HisTrap column (GE Healthcare) and finally eluted with 220 mM imidazole in buffer A. Eluate fractions were supplemented with 1 mM EDTA, 0.2 mM PMSF and 0.01% Tween-20, incubated overnight with Strep-Tactin Superflow high capacity resin (2-1208-010, IBA), and eluted with buffer B (50 mM TRIS-HCl pH 7.5, 200 mM NaCl, 5% (v/v) glycerol, 1 mM DTT) containing 10 mM desthiobiotin. After concentrating the eluate by ultrafiltration, final purification proceeded by size-exclusion chromatography with a Superose 6 column (GE Healthcare) pre-equilibrated in buffer B containing 1 mM MgCl2. Purified protein was snap-frozen and stored at −80°C until use. Most of the time, however, we used fresh condensin that had not been frozen.

### ATPase assay

We used a colorimetric phosphate detection assay (Innova Biosciences) to measure the ATPase activity of condensin complex. We mixed 50 nM condensin complexs and 50 ng/uL labmda DNA (Promega) for 15 mins in 40 mM Tris-HCl (pH7.5), 50 mM NaCl, 2.5 mM MgCl2, 2 mM DTT, 5 mM ATP. After 15 min incubation at room temperature, the released phosphate’s concentration was measured following the protocol of the manufacturer.

### Sample preparation for dry AFM

For the condensin-only sample preparation, we deposited 5 nM condensin onto newly cleaved mica surface (or 0.00001 % polylysine pretreated mica in Figure S3) for various conditions, namely with/without ATP and for EQ mutant condensin holocomplex with ATP in B1 buffer (20 mM Tris, 50 mM NaCl, 2.5 mM DTT, and 2.5 mM MgCl_2_). We used a 2.5 mM ATP concentration for WT/ATP or EQ/ATP conditions, and incubated the mixtures for 10 min. After depositing the sample for 20 seconds, we rinsed the mica using H_2_O, and dried the sample by using a gentle stream of N_2_. Then, we classified the structures into O-, I-, B-shapes, or kleisin-or hinge-opened states.

For the study on DNA and condensin interactions, we mixed 3 ng/μL of lambda DNA (D1501, Promega) and 5 nM of condensin in an Eppendorf tube for a 10 min incubation to induce condensin-DNA interaction. We then added 1 mM ATP and incubated the sample for an additional minute. We deposited the samples onto 0.00001 % (w/v) polylysine-treated mica for 20 seconds and rinsed the surface using 3 mL MilliQ water. Finally, we dried the sample using N_2_ gas.

To induce DNA loop formation by condensin, we incubated 3 ng/μL of lambda DNA with 1.5 nM condensin for 5 min in B1 buffer, and added 100 μM of ATP to this solution. We then incubated the mixture for 4 min. We deposited the sample (volume of 7 μL) onto the polylysine-treated mica for 20 sec, and rinsed the sample using 1 mL of buffer containing 50 mM MgCl_2_ to completely immobilize the DNA on the surface and prevent DNA diffusion during the washing and drying step. Then, the sample was rinsed with 1 to 3 mL MilliQ water. While rinsing the sample on the mica, we tilted the mica with a small angle (~10° with respect to horizontal direction) in order to reduce tension on the DNA sample. The sample was dried by a gentle stream of N_2_.

Upon using high polylysine concentrations, DNA is strongly attached on the surface as soon as it deposited, which yields kinetic trapping (Rivetti et al., 1996). The effective persistence length for such a projection is smaller than for an equilibrated structure that is obtained for DNA on MgCl_2_-treated mica. To prevent a high density of kinetically trapped DNA loops, we minimized the polylysine concentration to 0.00001 % (w/v), so that DNA was fixed onto the surface only intermittently using sparsely distributed polylysine molecules.

### Dry AFM imaging

We performed our AFM measurements in air on a Bruker Multimode AFM, with a Nanoscope V controller and Nanoscope version 9.2 software. We used Bruker ScanAsyst-Air-HR cantilevers (nominal stiffness and tip radius 0.4 N/m, and 2 nm, respectively) or Bruker Peakforce-HIRS-F-A (0.35 N/m and 1 nm). The imaging mode we used was PeakForce Tapping, with an 8 kHz oscillation frequency, and a peak force setpoint value less than 100 pN. For imaging the condensin structures, 3 μm × 3 μm scan areas with 2,048 × 2,048 pixels were recorded at 0.5 Hz scanning speed. To record condensin on DNA loops, images of 10 μm × 10 μm were imaged with 5,120 × 5,120 pixels at 0.2 to 0.7 Hz scanning speed. All measurements were performed at room temperature.

### Liquid AFM imaging

Freshly purified condensin holocomplex (2 nM) that had not been frozen was deposited on freshly cleaved mica surface with an imaging buffer (20 mM Tris [pH7.5], 50 mM NaCl, 1 mM ATP, 2.5 mM MgCl_2_, 2.5 mM DTT). After 10 s, the sample surface was rinsed with the imaging buffer. During the rinsing step, the sample was not dried. Following a published protocol (Uchihashi et al., 2012), we imaged the sample with HS-AFM (HS-AFM 1.0, RIB) using Nanoworld SD-S-USC-f1.2-k0.15 (2 nm tip radius, k = 0.15 N/m, f = 1.2 MHz) or Nanoworld SD-S-USC-f1.5-k0.6 cantilevers (2 nm tip radius, k = 0.6 N/m, f = 1.5 MHz). Typically, a scan size of 100 nm × 100 nm and 150 scan lines were used, with 2-10 Hz frame rates. For minimizing and stabilizing the sample-tip interaction during imaging, we used a feedback mechanism on the second harmonic amplitude (Schiener et al., 2004). We reconstructed movie files and images using a Matlab script. Using a Matlab code, we measured the hinge-head distance of every frame by measuring the distance between the two centers of mass of hinge and head domains. The hinge angle was measured as the angle between the two SMC arms at the hinge domain.

### Image processing

Before quantitative analysis, images were processed to remove background and transient noise data that would give false signals in automated analysis. This was done using Gwyddion version 2.53 (Nečas and Klapetek, 2012). To ensure that only the empty surface was used for background subtraction, an iterative procedure of masking particles and subtracting (planar and/or line-by-line) background polynomials was employed. Horizontal scars, which occur from time to time due to feedback instabilities or particles sticking to the AFM tip, were selected and removed by Laplacian background substitution. Finally, we used the blind tip estimation and surface reconstruction algorithms in Gwyddion to reduce the effects of AFM tip convolution (the widening of features due to the finite size of the AFM tip) on our images. It has been shown that the tip shape can affect estimates of SMC protein volume (Fuentes-Perez et al., 2012). Although blind tip estimation is not capable of fully correcting for the influence of tip shape, we found that it significantly improved the consistency of the volume estimates, and the increased contrast in the reconstructed images allowed for a better delineation of individual molecules in automated image quantification.

### Characterizing the condensin complex from dry AFM images

The volume of condensin molecules was measured using a home-built Matlab code. Background noise was subtracted by the program, and both the area and mean height of the structure was measured with respect to the background surface. The volume was then calculated by multiplying the deduced area and height. The distributions of the volumes were confirmed by Gywddion. To measure the volume using Gywddion, a height mask was applied to cover the condensin and grain measurement was used in order to determine the volume for the different masked objects.

Distances between the hinge-nonSMC subunits were measured between the hinge and nonSMC/head globular parts, using the line tool and measurement function of image J. The hinge angle was determined by measuring the angle between two SMC arms at the hinge domain.

### Measurement of DNA length and DNA loop size

We wrote Matlab scripts that perform automated image analysis to deduce the path of the DNA in images. The following procedure was used to select pixels that belong to areas covered with DNA: The image was smoothed with a Gaussian filter (*σ* = 0.5 pixels, window size = 5 pixels), and pixels with a height of more than 0.15 nm were selected. To get rid of noise and contaminations, objects (contiguous selected areas) with a maximum height less than 0.3 nm were removed from the selection, as well as objects of area less than 1000 nm^2^. To selectively analyze DNA on the surface, we excluded DNA-unbound proteins on the background by excluding objects with boundary length-to-area ratio > 0.1 nm^−1^ and those with area-to-bounding-box-area ratio < 0.25. As there was virtually no difference in observed height between the SMC arms and the DNA, condensin molecules touching the DNA were mostly included in the selection, and the total DNA length estimate is therefore a slight overestimate. The objects selected in this way were identified to be DNA, and visual inspection confirmed that this was reliable. To subsequently obtain the DNA length, the objects were skeletonized and the length was approximated as *d*_*p*_*n*_*s*_, with *n*_*s*_ the number of pixels of the skeleton, and *d*_*p*_ the linear size of a single image pixel. For measuring the size of individual loops, loops were identified manually and subsequently traced using DNA Trace software (Japaridze et al., 2017b).

### Estimation of the probability of accidental co-localization of condensin and DNA

The binding probability between condensin and DNA was estimated from AFM images as the fraction of proteins that overlapped with a DNA molecule. There is of course also a probability that a randomly deposited protein will overlap with DNA. This accidental co-localization probability is estimated as follows: We approximate the protein by its circumcircle with diameter *D_p_*. To overlap with the DNA, the center of this circumcircle should be within a distance *D_p_*/2 from the DNA. With a random placement of a condensin complex to the surface, the probability of this occurring in co-localization with the DNA is equal to the relative area of the image occupied by the space between two curves parallel to the DNA with a distance *D_p_* between them. If the total length of all DNA in the image is *L_DNA_*, this area can be approximated by *D_p_*×*L*_DNA_. See Figure S7A for a graphical illustration of this. Therefore, the probability of finding a randomly placed molecule co-localized with the DNA is

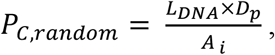

where *A*_*i*_ is the imaged area. We used a value of 40 nm for *D_p_*, and with an average *L*_DNA_ = 209 μm DNA in a 10×10 μm^2^ image, the accidental colocalization probability is about 8%. Note that this is an overestimate, since the condensin complex is often elongated so that in many cases the circumcircle can touch the DNA but the protein does not, and furthermore the DNA is curved and as a result the area between the parallel curves is less than *D_p_*×*L_DNA_*.

A similar line of reasoning can be applied to the localization of condensin at the loop stem (Figures S8A-S8C). Long DNA molecules that are attached to a surface have loops in them even in the absence of protein, simply due to the deposition of the randomly coiled DNA to the surface. There is a small chance that a randomly deposited protein, that was not already bound to the DNA before, locates with a distance *D_p_* from the loop crossing – a probability that can be calculated similar to above. Furthermore, a condensin that was already bound to the DNA before it got deposited, can, by chance, get located at the loop crossing in the process of depositing the DNA to the surface. Combining these two terms leads to an expression for the probability that condensin is accidentally observed as co-localized with loop crossings, which equals the number of DNA-bound condensins that are randomly located at the stem of DNA loops divided by the total number of DNA-bound condensins,

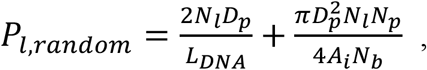

where *N_l_*, *N_P_* and *N_b_* are the total number of loops, total number of proteins, and the number of DNA-bound proteins in the image, respectively. For our typical conditions, this probability *P*_*l,random*_ is estimated to be about 3.5 %. See also Figure S8B.

### Width distributions of DNA-loop neck and protein

The neck size of the DNA was measured using the height profile tool of image J. We used the cross-section of the DNA region closest to the loop neck that did not overlap with the site of the protein. The DNA height profiles of the cross-section region showed two peaks, one from each side of the loop forming the neck, to each of which we fitted a Gaussian. The neck size was taken as the horizontal distance between the outer points of the FWHM of these Gaussians. The distance was defined this way as it gives a consistent definition, both when the DNA molecules or parts of the condensin complex were or were not resolved as separate Gaussian peaks. The same procedure was followed for measurements of the width of the protein that was located at the neck site.

### Loop extrusion visualization using single-molecule fluorescence microscopy

For the loop extrusion assay, we followed the protocol published by (Ganji et al., 2018).

