## Supplementary figures for "AFM images of open and collapsed states of yeast condensin suggest a scrunching model for DNA loop extrusion"

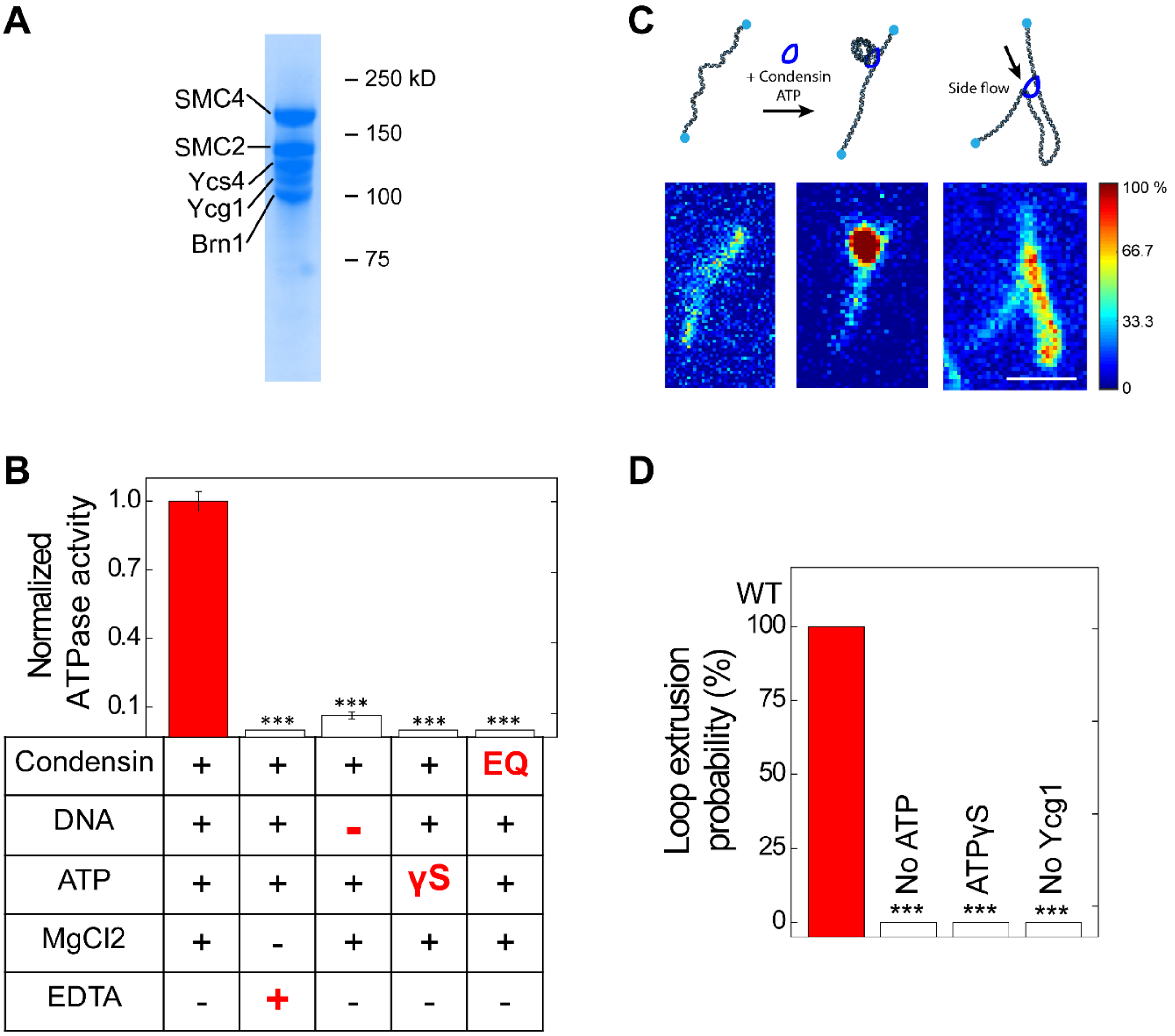


**Figure S1. Characterization of the functionality of yeast condensin.**

(A) SDS page gel of the condensin proteins. (B) ATPase activity of condensin with various controls (*N* = 3). (C) Loop extrusion assay with double tethered DNA showing that the condensin leads to the extrusion of a loop of DNA. (D) Loop-extrusion probability with various negative controls (*N* > 200).

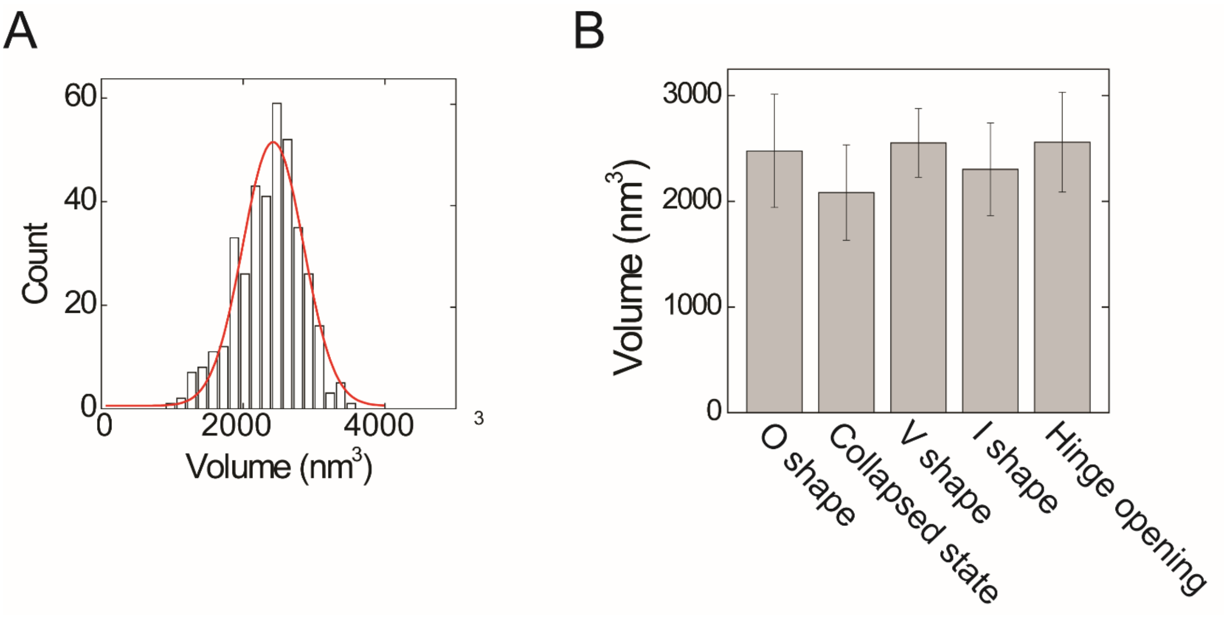


**Figure S2. Volume of the condensin holocomplex.**

(A) Frequency distribution of the condensin volume (*N* = 384) as measured from dry AFM images. Gaussian fitting yielded a mean value of 2420 ± 430 $nm^3$. (B) Mean volume of each shape conformation (*N* = 86, 200, 36, 51, and 11 for each state, respectively). There were no significant differences in molecular volume between the states, consistent with the fact that each contained the same subunits.

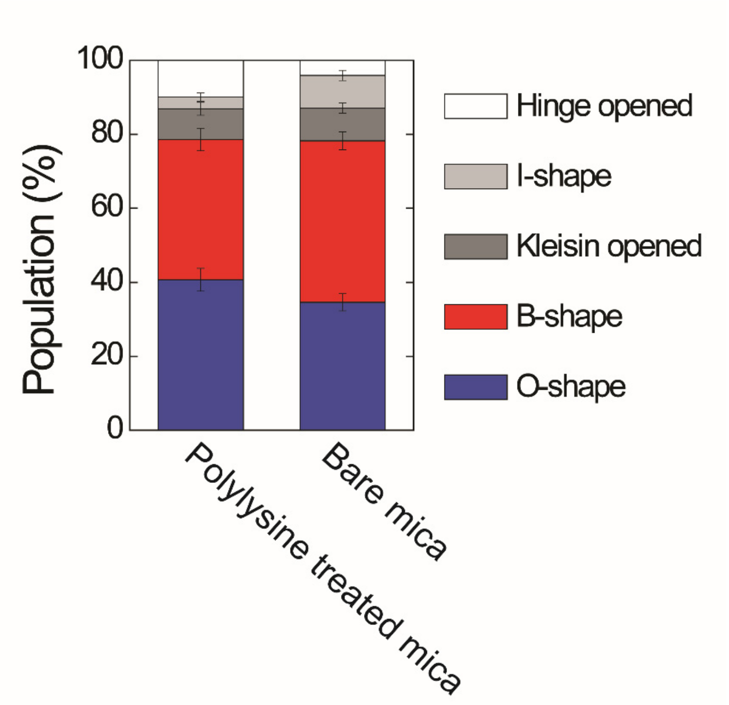


**Figure S3. Relative occurrence of different shapes of the condensin complex.**
Relative occurrence of different shapes of condensin in AFM images taken in the presence of ATP on polylysine-treated mica versus images on bare mica. Before sample deposition, polylysine was treated for 3 min, and the sample was rinsed using distilled water.

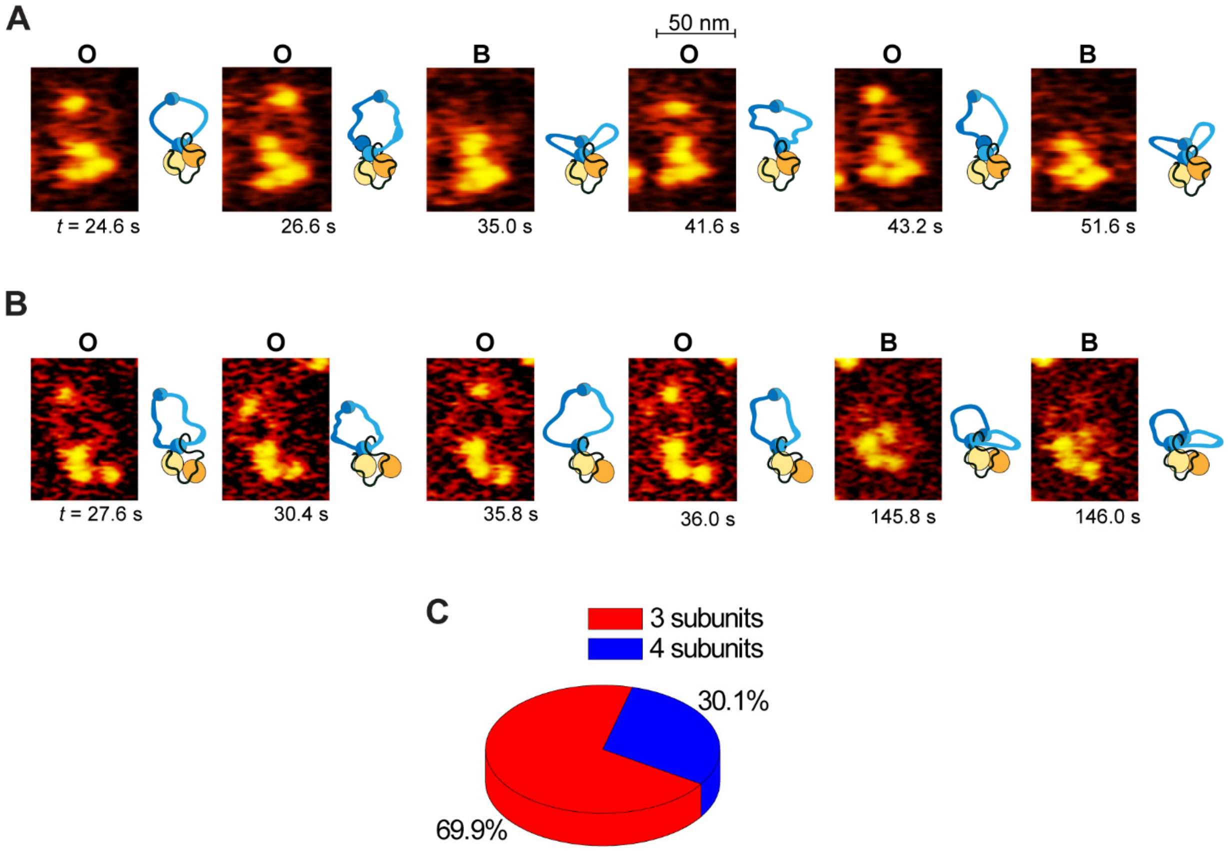


**Figure S4. Representative HS AFM image sequences of the condensin holocomplex.**
Images were taking from movies acquired at 5 frames/s rate, from Movie S2 (A) and Movie S3 (B). Changes between O- and B-shapes were observed. O and B indicate O-shape and B-shape, respectively. Cartoons on the right of each image are for visual guidance. (C) Number of distinguishable subunits in the globular domain (i.e., excluding the hinge) ($N > 4{,}000$ frames from 6 movies). About 70% of the complexes featured 3 distinguishable subunits in the globular domain, and about 30% featured 4 subunits. The complex with 3 subunits likely is composed of a dimerized head domain and two heat-domains, while two head domains were resolved separately in the case of the complex with 4 subunits in the globular domain.

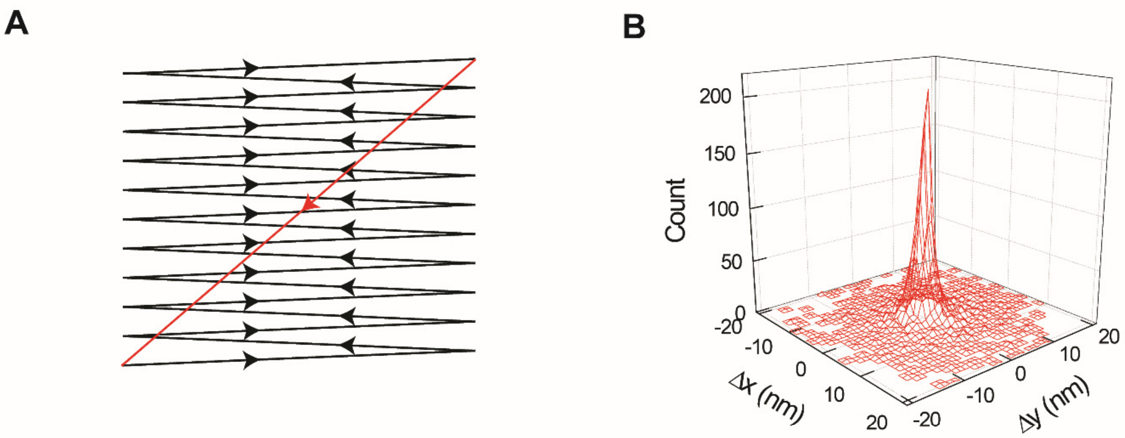


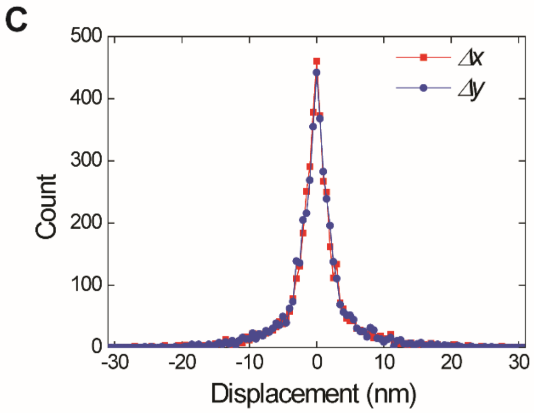


D

| | Δx | Δy |
|---|---|---|
| Standard deviation (nm) | 6.2 | 5.6 |
| Skewness (nm) | 0.2 | -0.14 |

**Figure S5. Tip scanning direction does not bias the hinge motion.**

In HS-AFM imaging in liquid, tapping and scanning the sample with the tip can potentially affect experiments adversely, e.g. induce a movement due to the tip pushing the molecule. The exact magnitude of the tip-sample forces in fluid tapping mode AFM is difficult to assess during imaging, but nevertheless can be qualitatively estimated from the magnitude of the second harmonic of the tapping signal (Schiener et al., 2004). We precisely controlled this and kept it to the minimum that still allowed for sufficient imaging resolution. To assess whether this was sufficient to avoid influencing the molecule dynamics, we analyzed the distribution of frame-to-frame position changes of the hinge domain with respect to the head domains. Since the scanning motion is not isotropic, one might expect an uneven distribution of hinge positional changes in the fast and slow scanning directions. Instead, we find a highly isotropic distribution, indicating minimal tip-induced effects. (A) The zigzag scanning direction of HS-AFM imaging. The horizontal direction scanning motion (black lines) is much faster than the vertical scanning. After finishing the imaging of a frame, the tip moves back to the starting point (red line) and scanning restarts. (B) Surface plot of the two-dimensional histograms of the hinge domain displacement at all time points ($N$ > 4,000 frames from 6 movies on different molecules). A radially symmetric distribution is obtained. No bias is observed towards a certain direction, also not towards the horizontal scanning direction (X-axis). (C) The distributions of X, Y displacements between adjacent images are nearly identical. (D) Tables of standard deviation and skewness of the X, Y displacements. Almost no skewness was observed in both cases.

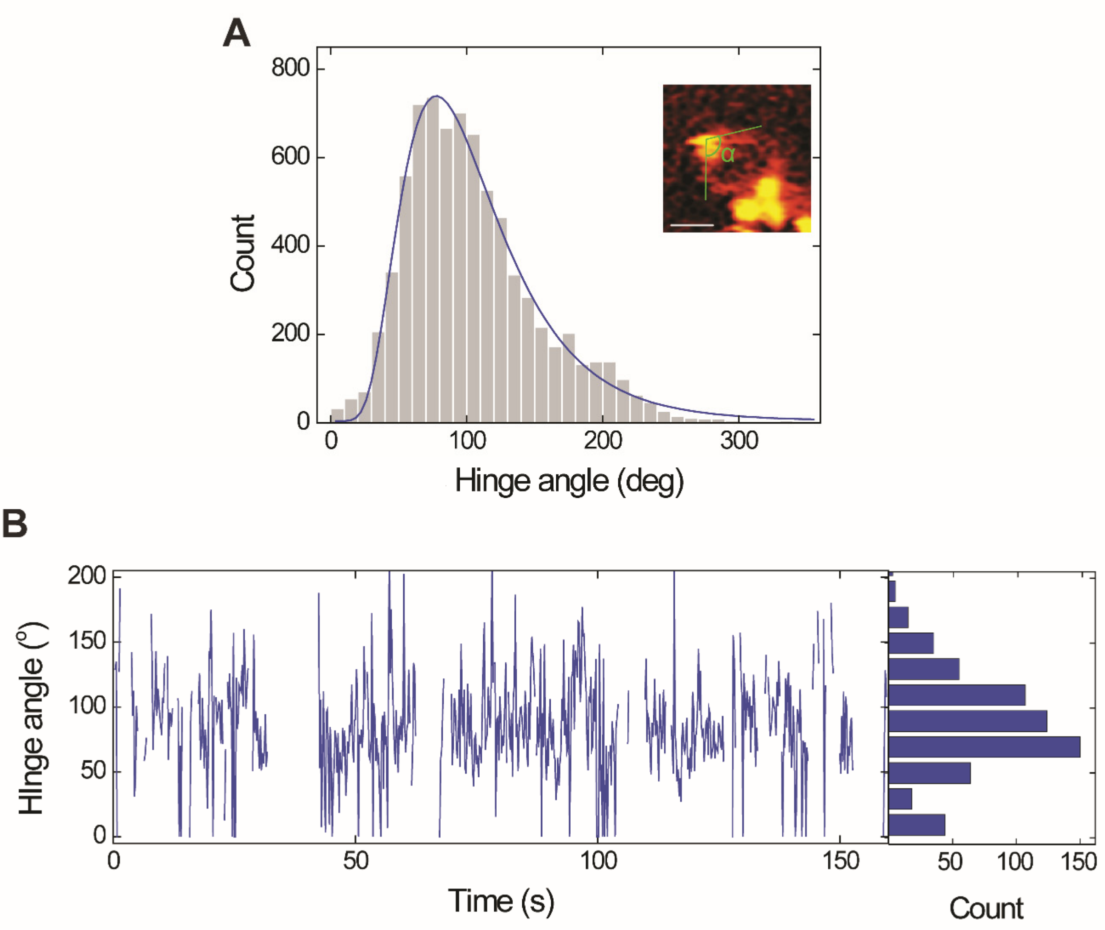


**Figure S6. Hinge angle – the angle between two SMC arms at the hinge domain.**

(A) Hinge angle distribution ($N$ > 4,000 frames from 6 independent molecules). The hinge angle was defined by the angle between two tangential lines of SMC arms at the hinge domain, see inset. Fit is a lognormal distribution with a mean of 96° and a standard deviation of 48°. (B) Representative trace of the hinge angle versus time. Right panel shows the distribution of the hinge angle from this trace.

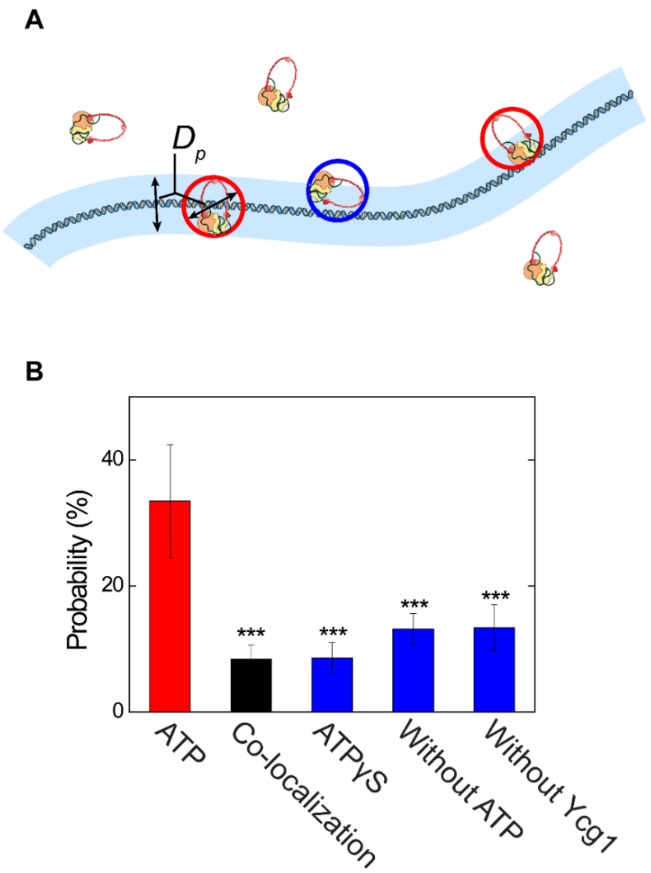


**Figure S7. Binding probability and co-localization probability of condensin on DNA.**

(A) Schematic that illustrates the calculation of the probability for DNA and condensin to co-localize. Each condensin is approximated by a circumcircle with size $D_p$ ~ 40 nm. The light blue shaded region indicates the area that should contain the center of the circumcircles for a randomly deposited condensin to co-localize with DNA. Condensin complexes inside the red circles overlap with DNA and would be counted as being bound in our measurements. Condensin complexes such as that in the blue circle are also included in the calculated co-localizing fraction, since their circumcircle overlaps with the DNA, even though the molecule itself does not touch the DNA. We deliberately err on the side of caution and do not correct for this, and the estimated value for the colocalization probability is therefore an upper bound. (B) The DNA binding probability of condensin with ATP (red bar), the (inadvertent) colocalization probability of condensin (black bar), and DNA binding probability of condensin for negative controls (blue bars) (N ≥ 5 independent samples). The DNA binding probability of condensin is the observed binding probability, while the colocalization probability is the estimated probability that DNA and condensin accidently overlap on an AFM surface without biochemical interactions.

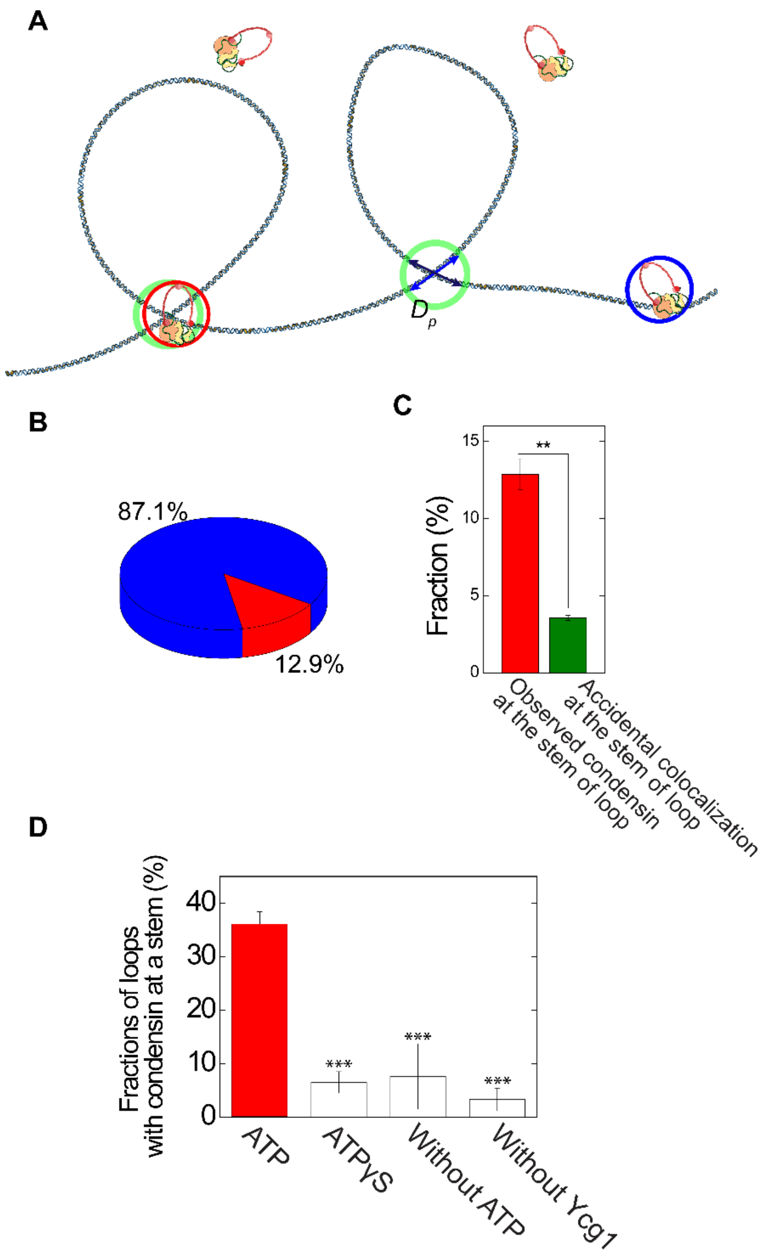


**Figure S8. Occurrence of condensin at the stem of DNA loops and elsewhere along the DNA.** (A) Schematic to illustrate our estimate of the localization probability of condensin at the stem of the loop versus elsewhere on the DNA. Green circles denote the loop-stem area with a size $D_p$. Condensin occurring there (red circle) are counted 'at the loop stem', whereas all other DNA-bound condensin are counted 'not at the loop stem' (blue circle). (B) Experimentally measured fractions of DNA-bound condensins that are observed at a loop stem (red), and not at a loop stem (blue) (*N*

= 5). (C) Fractions of DNA-bound condensins that are observed at a DNA loop stem (red), and the estimated accidental localization of condensin at a stem of DNA loops (green; $N = 5$ independent samples) – see Material and Methods. Errors are SEM. (D) Experimentally deduced fraction of DNA loops with condensin localized at the stem of the DNA loop with various controls. During sample preparation, two DNA regions can randomly overlap to form a loop-like structure on the mica surface. The occurrence of such loops in the AFM images will not depend on condensin or ATP. By contrast, the number of loops that are extruded by condensin complexes will depend on ATP hydrolysis. This indeed is reflected in the data, as the fractions of the number of loops that feature a condensin at the stem (among all loops with or without condensin at the stem) is significantly large in the presence of ATP and small in the controls ($N \geq 5$ independent samples).

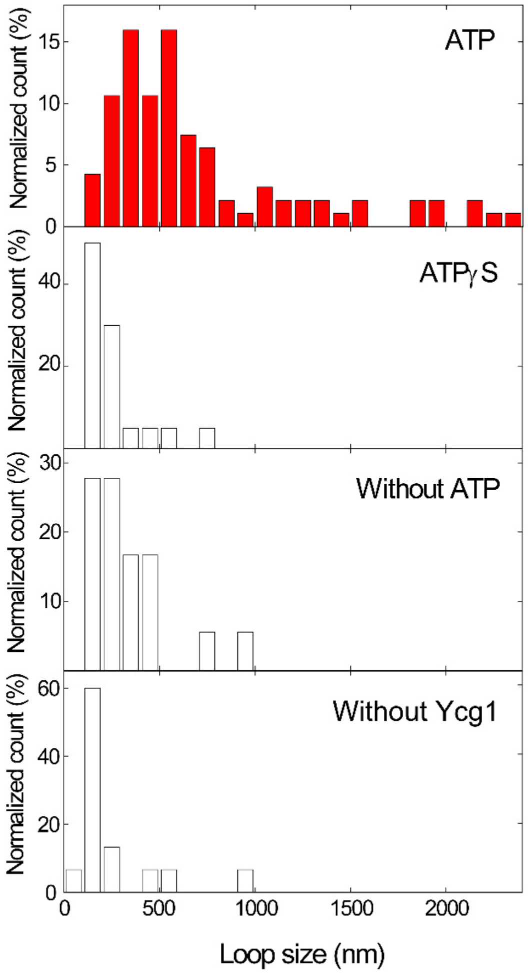


**Figure S9. Loop size distributions for various controls.**

DNA loop size distribution for data obtained with ATP, with ATPγS, , without ATP, and without Ycg1 (N = 94, 22, 20, and 11, respectively). The importance of these data lies in the larger loop size observed with ATP as compared to the data in the other three controls. The actual value of the median loop size (0.54 μm in top panel) is in fact rather dependent on experimental conditions (e.g. incubation time) and limited by our finite imaging area (10 μm × 10 μm) and DNA overlap, which hampers clean identification of large loops in our experiments. Hence, we measured relative small loop sizes (< 1 μm) compared to earlier fluorescence microscopy experiments (Ganji et al., 2018). In the data presented here, we measured the size of loops where condensin localized at the stem.

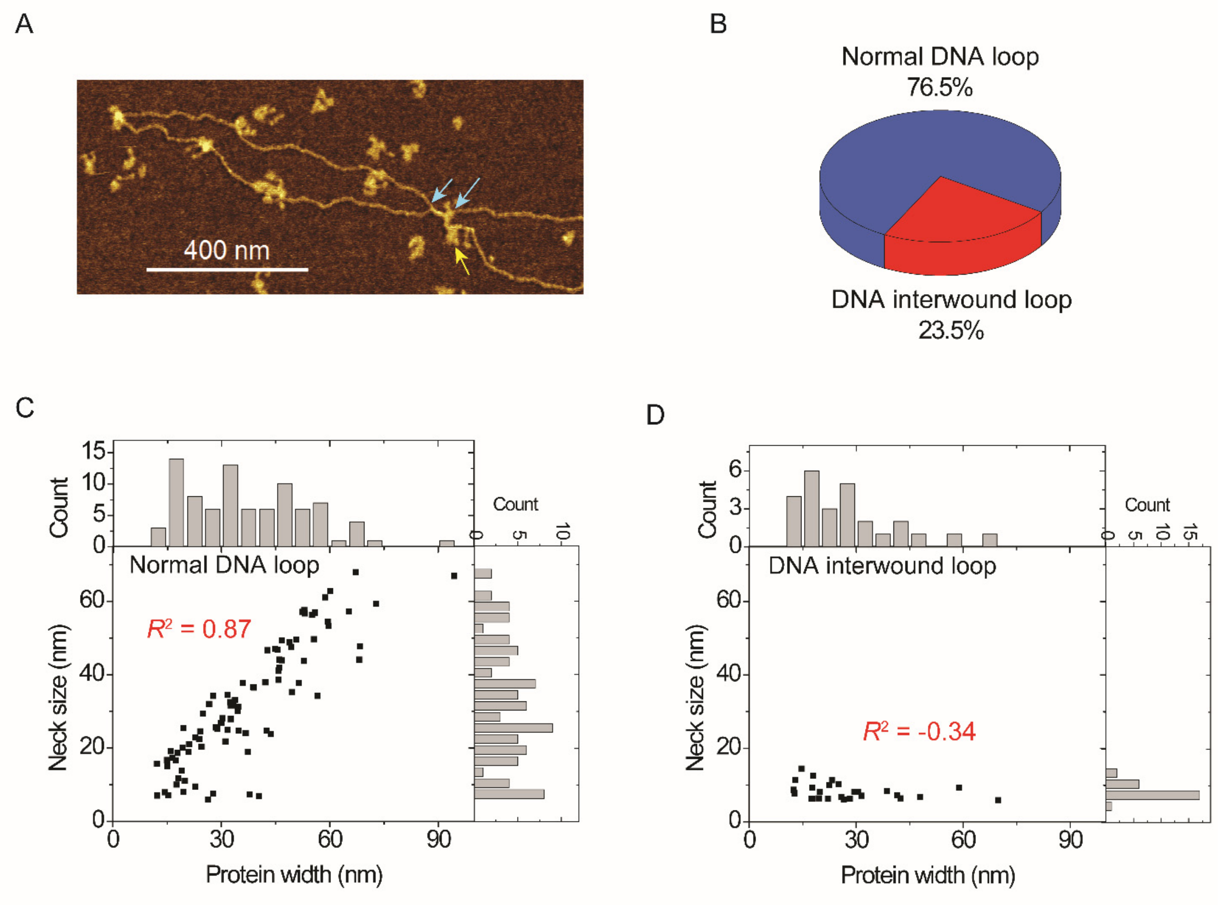


**Figure S10. Loops with an interwound DNA structure at the stem of loop.** (A) Example of an interwound DNA loop. Two arrows indicate the starting and ending points of an interwound DNA region. Yellow arrow indicates a condensin complex located near the loop stem. (B) Fraction of loops with an interwound region at the stem ($N$ = 116). (C) Scatter diagram of the protein width vs the neck size of normal (non-interwound) DNA loops. (D) Idem for DNA interwound loops at the stem. A neck size of ~9 nm is observed, which may be attributed to the two parallel DNA molecules (that feature a ~2 + 2 nm dsDNA width), convoluted by the AFM tip size.

**Supplementary Videos**

**Movie S1-S4: High-speed liquid AFM movies of condensin holocomplex, related to Figure 2 and S4.** Movie speed: 4x.

**Movie S5: Liquid AFM movie of condensin at the stem of DNA loop, related to Figure 4A.** Movie speed: 10x.
